# Modulation of polyamine metabolism in *Arabidopsis thaliana* by salicylic acid

**DOI:** 10.1101/2020.10.28.359752

**Authors:** Franco R. Rossi, Andrés Gárriz, María Marina, Fernando L. Pieckenstain

**Author notes:** Franco R. Rossi, Andrés Gárriz, María Marina, Fernando L. Pieckenstain. Corresponding author, Fernando L. Pieckenstain.

## Abstract

Polyamines (PAs) play important roles in plant defense against pathogens, but the regulation of PA metabolism by hormone-mediated defense signaling pathways has not been studied in depth. In this study, the modulation of PA metabolism in Arabidopsis by salicylic acid (SA) was analyzed, by combining the exogenous application of this hormone with the use of PA biosynthesis and SA synthesis/signaling mutants. SA induced notable modifications of polyamine metabolism, mainly consisting in putrescine accumulation both in whole-plant extracts and apoplastic fluids. Put was accumulated at the expense of increased biosynthesis by arginine decarboxylase 2 and decreased oxidation by copper amine oxidase. Enhancement of Put levels by SA was independent of the regulatory protein Non-Expressor of Pathogenesis Related 1 (NPR1) and the signaling kinases MKK4 and MPK3, but depended on MPK6. On its part, plant infection by *Pseudomonas syringae* pv. *tomato* DC3000 elicited Put accumulation in a SA-dependent way. The present study demonstrates a clear connection between SA signaling and plant PA metabolism in Arabidopsis and contributes to understand the mechanisms by which SA modulates PA levels during plant-pathogen interactions.

**Highlight:** Salicylic acid modulates polyamine biosynthesis and catabolism in Arabidopsis. Regulatory effects of salicylic acid are independent of the master regulator NPR1 and are mediated by the MPK6 kinase.

## Introduction

In nature, plants are continuously challenged by pathogenic microorganisms that pose serious threats to their growth and development. Nevertheless, these biological interactions rarely result in disease because plants have developed, through their evolution, sophisticated defense mechanisms that allow them to avoid the deleterious consequences of infection (Berens *et al*., 2017). In this trend, after the recognition of the pathogen, plants activate different local and systemic signaling pathways mediated by hormones such as salicylates, jasmonates and ethylene, with the purpose to hinder the propagation of the microorganism to the rest of the plant and/or prepare the plant for future attacks. These responses involve the activation of protein kinases conducing to the expression of defense proteins and the accumulation of different compounds that globally contribute to plant defense.

Salicylic acid (SA) is a plant defense hormone that plays critical roles in plant immunity. Signaling cascades mediated by this hormone are required for the activation of key processes involved in plant defense. Thus, SA accumulation is associated with the hypersensitive response, a type of programmed cell death usually induced during effector-triggered immunity (Radojičić *et al*., 2018). In addition, SA participates in the establishment of the systemic acquired resistance, a set of defense responses triggered in uninfected distal tissues that helps plants to defend themselves against further infections by a broad spectrum of pathogens. In *Arabidopsis*, downstream SA signaling is mainly coordinated via the transcriptional regulator NPR1, whose degradation is controlled by its paralogues NPR3/NPR4 in a SA concentration-dependent manner (Fu *et al*., 2012; Moreau *et al*., 2012), and specific MAPK (also named MPK) proteins that exert positive (AtMPK3 and AtMPK6) and negative (AtMPK4) effects on defense gene expression (Colcombet and Hirt, 2008). Additionally, SA was shown to activate defense responses by an NPR1-independent pathway (Uquillas *et al*., 2004; Herrera-Vásquez *et al*., 2015; Singh *et al*., 2018).

PAs, mainly the diamine putrescine (Put), the triamine spermidine (Spd) and the tetraamine spermine (Spm), are natural aliphatic polycations essential for prokaryotic and eukaryotic cells. As PAs are positively charged at physiological pH, they interact with anionic molecules such as proteins, nucleic acids and phospholipids. As a consequence, they modulate protein–protein (Thomas *et al*., 1999; Garufi *et al*., 2007) and DNA–protein interactions (Shah *et al*., 1999; D’Agostino *et al*., 2005), as well as RNA structure (Igarashi and Kashiwagi, 2010). In this way, PAs regulate many fundamental cellular processes such as cell division, differentiation and proliferation, cell death, DNA and protein synthesis, gene expression, and stress responses (Seiler and Raul, 2005; Kusano *et al*., 2008; Igarashi and Kashiwagi, 2010; Romero *et al*., 2018). In plants, different routes are involved in the synthesis of Put. Thus, this PA may be obtained after the decarboxylation of ornithine by ornithine decarboxylase (ODC). On the other hand, arginine can be converted into agmatine by the enzyme arginine decarboxylase (ADC), which is later metabolized to Put in two additional reactions catalyzed by agmatine iminohydrolase and N-carbamoylputrescine amidohydrolase. A novel route for Put production from arginine was recently described to operate in Arabidopsis and soybean. This pathway is located in chloroplasts and involves arginases that also deploy agmatinase activity (Patel *et al*., 2017). The synthesis of the higher PAs Spd and Spm requires decarboxylated *S*-adenosylmethionine generated by the action of *S*-adenosylmethionine decarboxylase (SAMDC). Decarboxylated *S*-adenosylmethionine functions as an aminopropyl donor in the reactions catalyzed by spermidine synthase (SPDS) and spermine synthase (SPMS), which in two consecutive reactions add aminopropyl moieties to convert Put into Spd, and Spd into Spm, respectively. Interestingly, Arabidopsis has lost the ODC gene and thus depends exclusively on the arginine routes for Put synthesis (Hanfrey *et al*., 2001; Fuell *et al*., 2010). As a consequence, modulation of ADC activity in this species is of key importance for the regulation of PA levels. Arabidopsis possess two ADC genes (*AtADC1* and *2*), which are differentially regulated during ontogenesis (Hummel *et al*., 2004) and in response to distinct environmental stimuli (Alcazar *et al*., 2012; Rossi *et al*., 2015). Simultaneous knockout of both *AtADC* genes leads to a lethal phenotype, thus confirming that *Arabidopsis* strongly depends on ADC for Put biosynthesis (Urano *et al*., 2005). In addition to *de novo* biosynthesis, PA levels are also regulated by catabolic pathways. PAs are oxidatively deaminated by amine oxidases (AOs), which can be classified into FAD-dependent (PAOs) and copper-containing (CuAO) enzymes. PAOs catalyze the oxidative deamination of Spm, Spd and/or their acetylated derivatives at the secondary amino group, whereas CuAOs catalyze the oxidation of Put at the primary amino group (Planas-Portell *et al*., 2013). Plant AOs contribute to important physiological processes, not only through the regulation of cellular PA levels, but also through their reaction products: aminoaldehydes, gamma aminobutyric acid and in particular, H_2_O_2_ (Cona *et al*., 2006).

A considerable amount of information about the role played by PAs during the interactions of plants with pathogenic microorganisms is available. Thus, significant changes in the expression of plant genes encoding PA biosynthetic and catabolic enzymes have been reported, which are usually associated to variations in the concentration of these compounds in plant tissues (Jiménez-Bremont *et al*., 2014). For instance, PA accumulation by the induction of *de novo* biosynthesis, followed by further PA oxidation by AOs were found to play an important role in plant defense against pathogens (Marina *et al*., 2008; Moschou *et al*., 2009; Gonzalez *et al*., 2011). A great progress has been made in the last years towards a better understanding of the physiological significance of these changes. Thus, the expression of both ADC isoforms was reported to increase during the hypersensitive response triggered in *Arabidopsis* by avirulent cucumber mosaic virus (Mitsuya *et al*., 2009), as well as during the infection of this plant by the cyst nematode *Heterodera schachtii* (Hewezi *et al*., 2010). Nevertheless, we later demonstrated that even though there is a certain degree of functional redundancy between the two ADC isoforms, *ADC2* has a major contribution to total ADC activity in *Arabidopsis*. In addition, our work demonstrated that only *ADC1* is induced in response to *Pseudomonas viridiflava* infection (Rossi *et al*., 2015). In this regard, it was reported that *Arabidopsis* MPK3 and MPK6 play a positive role in the regulation of Put biosynthesis, and that Put contributes to *Arabidopsis* defense against *P. syringae* (Kim *et al*., 2013). Besides, a role for Spm in modulating global changes in the expression of defense-related genes in *Arabidopsis* has also been described (Mitsuya *et al*., 2009; Sagor *et al*., 2009; Gonzalez *et al*., 2011).

Many reports have described clear connections between defense-related hormones and PA metabolism. For instance, treatment of barley and wheat plants with methyl-jasmonate leads to increased PA levels and activities of enzymes involved both in PA biosynthesis and oxidation (Walters *et al*., 2002; Haggag and Abd-El-Kareem, 2009). Additionally, treatment of *Arabidopsis* plants with methyl-jasmonate increases the expression of *ADC2* while *ADC1* remains unaltered, suggesting the existence of different regulatory pathways for both genes (Perez-Amador *et al*., 2002). In turn, treatment with SA induces PA accumulation by activating ADC and ODC expression in maize, tobacco, and tomato (Németh *et al*., 2002; Jang *et al*., 2009; Zhang *et al*., 2011). In chickpea plants, however, the application of SA repress the induction of JA-mediated PA oxidation (Rea *et al*., 2002). Recently, Liu et al (2020) found that defense signaling triggered by Put partly depends on SA accumulation. Moreover, this study demonstrated that Put induces local SA accumulation, as well as local and systemic reprogramming of genes involved in systemic acquired resistance. Therefore, a number of evidences demonstrate the existence of a cross-talk between SA and PA metabolism. However, current knowledge about the mechanisms by which SA regulates PA metabolism during plant infection by pathogens is far from being complete. In the present work we demonstrate that the exogenous application of SA induces notable modifications of PA metabolism in *Arabidopsis*, mainly consisting in an increase of Put levels at the expense of ADC activity induction. These changes were found to be NPR1-independent and partially dependent on MPK6 activity. In addition, we also found that Put supplementation reduces plant susceptibility to *P. syringae*. Overall, these findings indicate a clear interconnection between SA signaling and plant PA metabolism and also contribute to understand the mechanisms by which SA modulates PA metabolism during plant-pathogen interactions.

## Materials and methods

### Plant material and growth conditions

The *Arabidopsis thaliana* Columbia-0 (Col-0) was used as wild-type and background ecotype when mutant or overexpressor plant lines were used. Plant lines *mpk3-2, mpk6-2* and *mkk4* were acquired from SALK (SALK_151594, SALK_073907 and SALK_058307, respectively), *npr1-1* was kindly provided by Dr. María Elena Álvarez. Seedlings were grown under axenic conditions, after surface-disinfection with 75% (v/v) ethanol for 1.5 min followed by 5% (v/v) commercial bleach for 15 min and thorough rinsing with sterile distilled water. Disinfected seeds were sown on 24-well plates, two seeds per well, containing 1.5 ml of MS medium (0.8 % agar) supplemented with 3% (w/v) sucrose and stratified at 4°C for 2 d in the dark. Additionally, some experiments were performed using Petri dishes containing MS medium as described before. Plates were subsequently incubated for 2 weeks in a growth chamber with a 16-h light/8-h dark photoperiod at 24/22 °C, 55/65% relative humidity (day/night) and a photon flux density of 100 μmol m^2^ sec^-1^ provided by cool-white and *Grolux* fluorescent lamps.

### SA treatment

SA supplementation was performed by adding 50 μl of a stock solution to each well in order to reach a final concentration of 50, 100 and 250 μM. SA stock solutions were added to 15-days old Arabidopsis by touching the wall of each well with a tip, avoiding the direct contact of the solution with the seedlings. No deleterious effects were evident in seedlings treated with SA under the conditions used in the present work. Additional information about the system set-up is provided in Figures S1 and S2.

### ADC and CuAO activity

*In vitro* ADC activity was measured by quantifying the ^14^CO_2_ released, employing [U-^14^C]-L-arginine as a substrate. Whole seedlings were grinded in a mortar with liquid nitrogen immediately after the harvest. The homogenates were centrifuged for 15 min at 10000 x g and the supernatants were used as enzyme source according to the method previously described by Rossi *et al*. (2015).

CuAO activity in apoplastic washing fluids (AWFs) was measured by quantifying the H2O2 released using the homovanillic acid oxidation method, as previously reported by Rodriguez *et al*. (2009). Proteins were precipitated from AWFs with a saturated solution of (NH_4_)_2_SO_4_ using a typical two-step protocol. The hydrogen peroxide released by 100 μl-samples was assessed by H_2_O_2_-dependent oxidation of homovanillic acid. Changes in the fluorescence of homovanillic acid (55 μM at 407 nm (em) and 305 nm (ex)), in the presence of 12 μg·ml^-1^ horseradish peroxidase and 1 mM Put, were measured after one hour of incubation at 37 °C using a multi-plate reader *BioTek Synergy H1*. In all cases, reactions were stopped before fluorescence determination by adding 5 μl 5N NaOH.

### Determination of free polyamine concentration

PA levels were determined as described by Rossi *et al*. (2015). PA were extracted by grinding 200 mg fresh plant material in 0.6 ml 5% (v/v) perchloric acid with a plastic pestle and incubating extracts overnight at 4 °C. After centrifugation at 10,000 x g for 15min, 6 μl of 0,1 mM 1,7-heptanediamine were added as internal standard to 60 μl aliquots of plant extracts. Then, 60 ml of saturated Na_2_CO_3_ and 75 μl of dansyl chloride (10 mg ml^-1^ acetone) were added and the mixture was incubated 2 h at 55 °C. Reaction was stopped by adding 25 μl of proline (100 mg ml^-1^) and dansylated PAs were extracted in 200 μl of toluene. The organic phase was vacuum-evaporated and dansylated PAs were dissolved in 40 μl of acetonitrile and analyzed by reversed phase HPLC using a *Waters 1525* binary HPLC pump and a *2475 Multi λ* fluorescence detector, as described previously (Marcé *et al*., 1995).

### Bacterial growth, plant inoculation and disease evaluation

*Pseudomonas syringae* pv. *tomato* strain DC3000 (*Pst*) was routinely grown in LB medium containing 50 μg ml^-1^ rifampicin at 28 °C. Inocula were prepared by growing the bacterium in LB medium overnight and bacterial cultures were centrifuged and washed twice just prior to inoculation. Finally, bacteria were suspended in sterile distilled water at indicated bacterial cell densities (600 nm).

A flood-inoculation method described by Ishiga *et al*. (2011) was used in the present work. Briefly, 40 ml-aliquots of a bacterial suspension in sterile distilled water containing 0.015% Silwet L-77 were dispensed in 24-well culture plates or Petri dishes containing 2-week-old Arabidopsis seedlings, as described in “Plant Material and growth conditions”, in order to flood them. Then, plates were incubated for 90 s at room temperature and the bacterial suspension was removed draining it in a beaker, remnants of bacterial solution were carefully removed by pipetting. Plates containing inoculated seedlings were sealed with Saran™ wrap and incubated as described in “Plant Material and growth conditions”. Control plants were treated in the same way with sterile distilled water containing 0.015% Silwet L-77.

Bacterial infection of inoculated plants was estimated on the basis of the number of colony forming units (CFUs) retrieved after surface-disinfection at different times after inoculation. For this purpose, aerial parts of plants were collected, weighed and disinfected with 5% commercial bleach for 3 min. After washing three times with sterile distilled water, plant material was homogenized in 0.4 ml sterile distilled water using a plastic pestle, and dilutions were plated on LB medium containing rifampicin 50 μg ml^-1^. Treatments comprised 4-5 replicates, each consisting on a pool of aerial parts collected from two seedlings. Plates were incubated for 24 h at 28 °C and CFUs were counted using properly diluted samples under a stereoscopic microscope.

### Gene expression analysis

Pools of 48 seedlings were frozen in liquid nitrogen and total RNA was extracted with TRI reagent (*Sigma Chemicals*) according to the manufacturer’s instructions. First-strand cDNAs were synthesised using Moloney Murine Leukemia Virus Reverse Transcriptase (*Promega*, www.promega.com). The quantification of gene expression was made by qRT-PCR, 1 μl of synthesised first-strand cDNA (1:5 dilution) was further diluted to 7.5 μl with sterile distilled water and the same volume of *FastStart Universal SYBR Green Master* (with Rox) was added to a final volume of 15 μl. Primers used in these reactions are listed in Table S1. Reactions were performed in a *StepOne Plus* qPCR system with the aid of the *StepOne Plus qPCR Software v2.3* (*Applied Biosystems*). Relative quantification was performed by the comparative cycle threshold method (Pfaffl, 2001) using *InfoStat Software v2017.1.2* (Di Rienzo *et al*., 2016).

### GUS activity

GUS activity was measured using a fluorescent assay. Whole seedlings (40-50 mg) were macerated with a plastic pestle in microfuge tubes containing 300 μl of base buffer (50mM Na_2_HPO_4_; 10 mM β-mercaptoethanol; 10 mM EDTA pH 8; sarcosyl 0.1%; Triton X-100 0,1%). Extracts were clarified by centrifugation at 4 °C, and aliquots were diluted 1/10 with base buffer. Enzymatic reactions were performed in a final volume of 500 μl in the presence of 1 mM of 4-methylumbelliferyl-beta-D-glucuronide (MUG) during 1 h at 37 °C. Finally, 100 μl of enzymatic reaction were stopped adding 900 μl of 0.2 M Na_2_CO_3_. Fluorescence was measured with excitation at 365 nm and emission at 455 nm in a multi-plate reader *BioTek Synergy H1* and calibration curves were made with 4-methylumbeliferone (4-MU). Total protein concentration was determined using Bradford’s method.

### AWFs extraction

The AWFs were obtained as described by O’Leary *et al*. (2014) with some modifications. Whole seedlings were carefully removed from 24-well plates containing MS medium, washed in distilled water and excess moisture was removed with absorbent paper. Seedlings were infiltrated with Tris-HCl buffer (100 mM, pH 6.5) creating negative pressure within a 50-ml syringe by pulling and releasing slowly the plunger. Then, seedlings were dried with absorbent paper and inserted into 10 ml-syringes, which were then introduced into 50-ml polyethylene tubes that were subsequently centrifuged at 1500 × g for 20 min at 4 °C. The AWFs recovered were subjected to additional centrifugation at 12,000 × g for 10 min to remove cells and particulate matter and were stored at −20 °C until use. Absence of cytosolic contamination was confirmed by the absence of glucose-6-phosphate dehydrogenase activity in the AWFs.

## RESULTS

### Selection of exogenous SA concentrations for induction of defense responses

As stated previously, the present work aimed to analyze the effects of SA on different aspects of PA metabolism in *Arabidopsis*. Some of the experiments hereby performed were based on the application of exogenous SA to Arabidopsis plants cultivated *in vitro*. Even though SA is naturally present in plants and plays a crucial role in the regulation of plant immunity, high levels of this compound can compromise defense signaling pathways controlled by other plant hormones and also inhibit plant growth and development (Koornneef and Pieterse, 2008; Berens *et al*., 2017). Therefore, in order to set up adequate experimental conditions we first examined the exogenous concentrations of SA capable to activate defense responses and protect *in vitro*-grown plants against *Pst*.

For this purpose, we analyzed the effects of SA on GUS activity in a transgenic *A. thaliana* line harboring the *PR1::GUS* reporter construction. *PR1* (pathogenesis-related protein 1) is a useful marker that reflects the activation of SA-mediated responses. As shown in Fig. S1, GUS activity was increased in plants treated with 50-500 μM SA. In addition, *Pst* multiplication was evaluated in WT Col-0 plants treated with SA prior to inoculation. In this case, a significant decrease in bacterial titers was detected in plants treated with 50 to 250 μM SA, as compared to plants inoculated without previous treatment with this hormone (Fig. S2). On the basis of these results, we decided to use 250 μM as the maximum concentration of SA in further experiments.

### Exogenous SA increases Put levels in whole-plant extracts and AWFs

We next determined whether SA modifies PA levels in Arabidopsis. The addition of SA to the culture medium of WT Col-0 plants caused a dose-dependent increase in the concentration of Put in whole plant tissues 24 and 48 hours post-treatment (HPT) (Fig. 1). Thus, plants supplemented with 100 and 250 μM SA respectively exhibited a 66% and 1-fold enhancement of Put concentration 24 HPT, as compared to control plants, (Fig. 1). This effect was magnified 48 HPT, and a significant increase in Put levels was also detected in plants treated with 50 μM SA. In turn, SA caused minor effects on the levels of the higher PAs Spd and Spm. Thus, a small increment in the concentration of Spd was evident for the highest SA concentration (250 μM) 48 HPT, whereas a small decrease in Spm levels was observed 24 HPT (Fig. 1).

**Figure 1.**
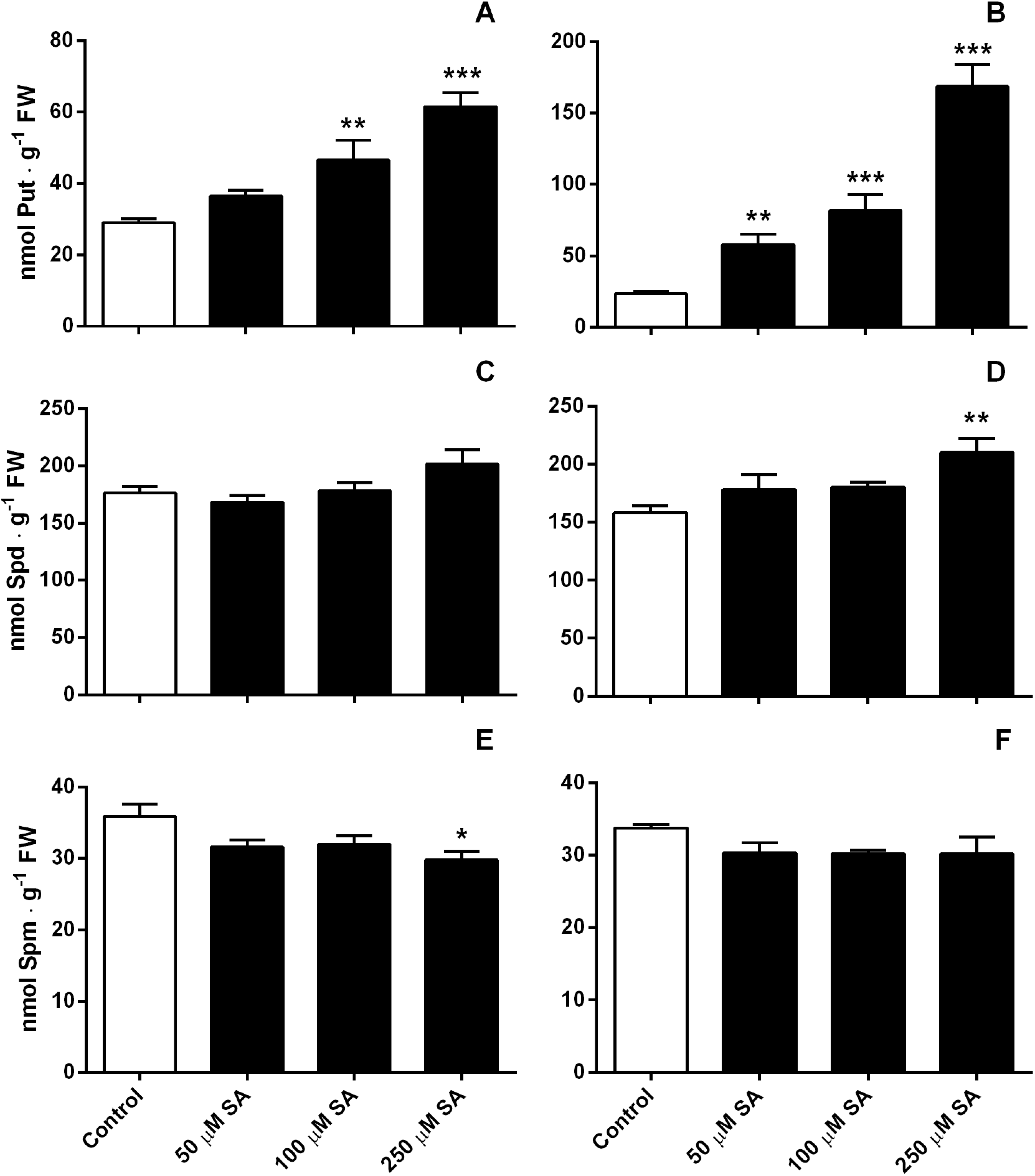
Free PA levels in extracts of WT Arabidopsis seedlings after treatment with SA. Fifteen-day-old *A. thaliana* seedlings were treated with several concentrations of SA (50, 100 and 250 μM) and samples were harvested at different times post treatment. PAs were extracted from control and SA-supplemented seedlings at 24 (**A-, C-, E-**) and 48 (**B-, D-, F-**) HPT. Free Put (**A-, B-**), Spd (**C-, D-**) and Spm (**E-, F-**) were determined by HPLC after derivatization with dansyl chloride. Results are means of five replicates ± SEM and statistically significant differences between SA-treated plants and controls are expressed as (*P*≤0.05), (*P*≤0.01) and (*P*≤0.001), according to oneway ANOVA and Dunnett’s Test.

Previous reports demonstrated that apoplastic PAs undergo significant changes during the establishment of pathogenic interactions, which play an important role in plant defense (Marina *et al*., 2008; Moschou *et al*., 2008; Vilas *et al*., 2018; Liu *et al*., 2019). This prompted us to evaluate the impact of SA in the contents of PAs in AWFs of *A. thaliana*. This analysis showed that the relative abundance of each PA in AWFs was different to whole-plant extracts. In this regard, Put levels found in AWFs were much closer to those of Spd than in whole plant extracts. The amendment of the culture medium with 250 μM SA caused a 70% increase in the levels of Put 48 HPT, whereas Spd and Spm concentration were reduced by 66% (Fig. 2).

**Figure 2.**
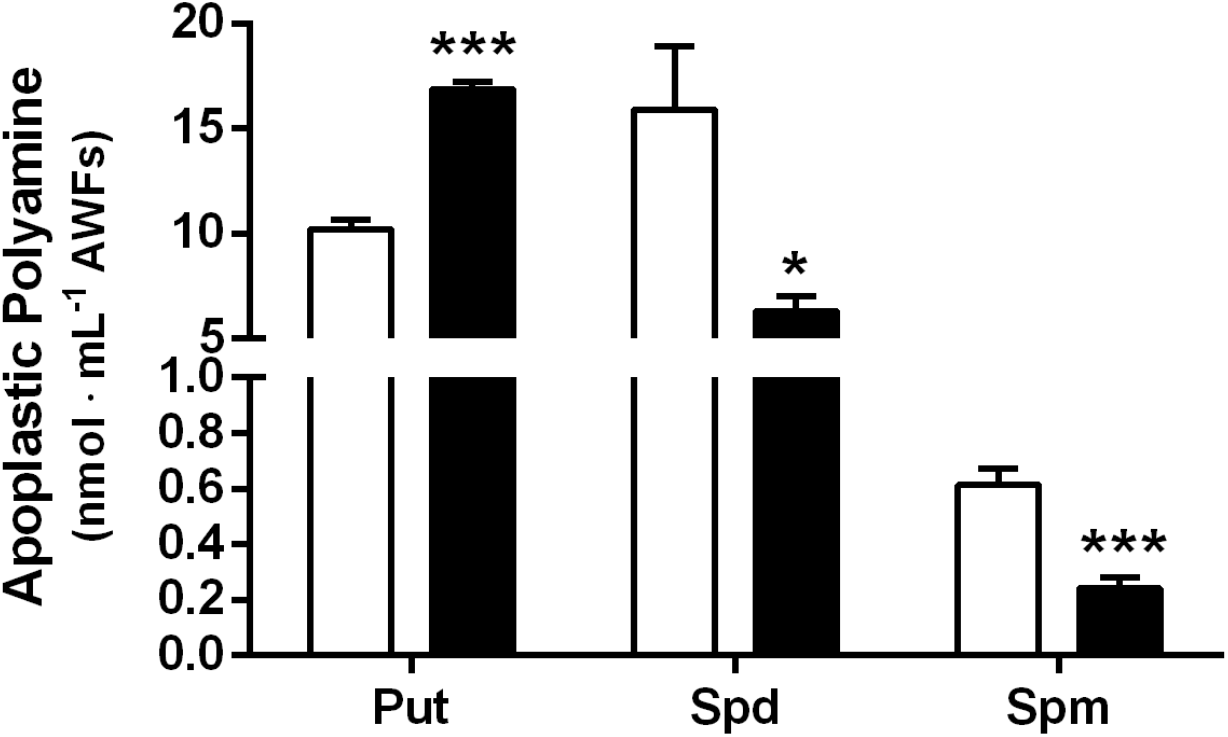
Free PA levels in apoplastic washing fluids (AWFs) of WT Arabidopsis seedlings. Free Put, Spd and Spm were extracted from AWFs of Arabidopsis seedlings supplemented with SA for 48 h. White and black bars represent controls and samples supplemented with SA (250 μM), respectively. Free PAs were determined by HPLC after derivatization with dansyl chloride. Results are means of five replicates ± SEM and statistically significant differences are expressed as *(*P*≤0.05) and ***(*P*≤0.001), according to Student’s t-test.

### Put increases Arabidopsis resistance to *Pst*

Taking into account that SA induced Put accumulation, we speculated that this PA might play a role in defense responses triggered by SA and in turn contribute to increase *A. thaliana* resistance to pathogen infection. In order to test this hypothesis, we assessed *Pst* multiplication in plants grown in the presence of 5 and 50 μM Put. In agreement with the abovementioned hypothesis, bacterial multiplication was reduced by 50% in Put-treated plants (Fig. 3), indicating that this diamine enhances plant resistance to *Pst*.

**Figure 3.**
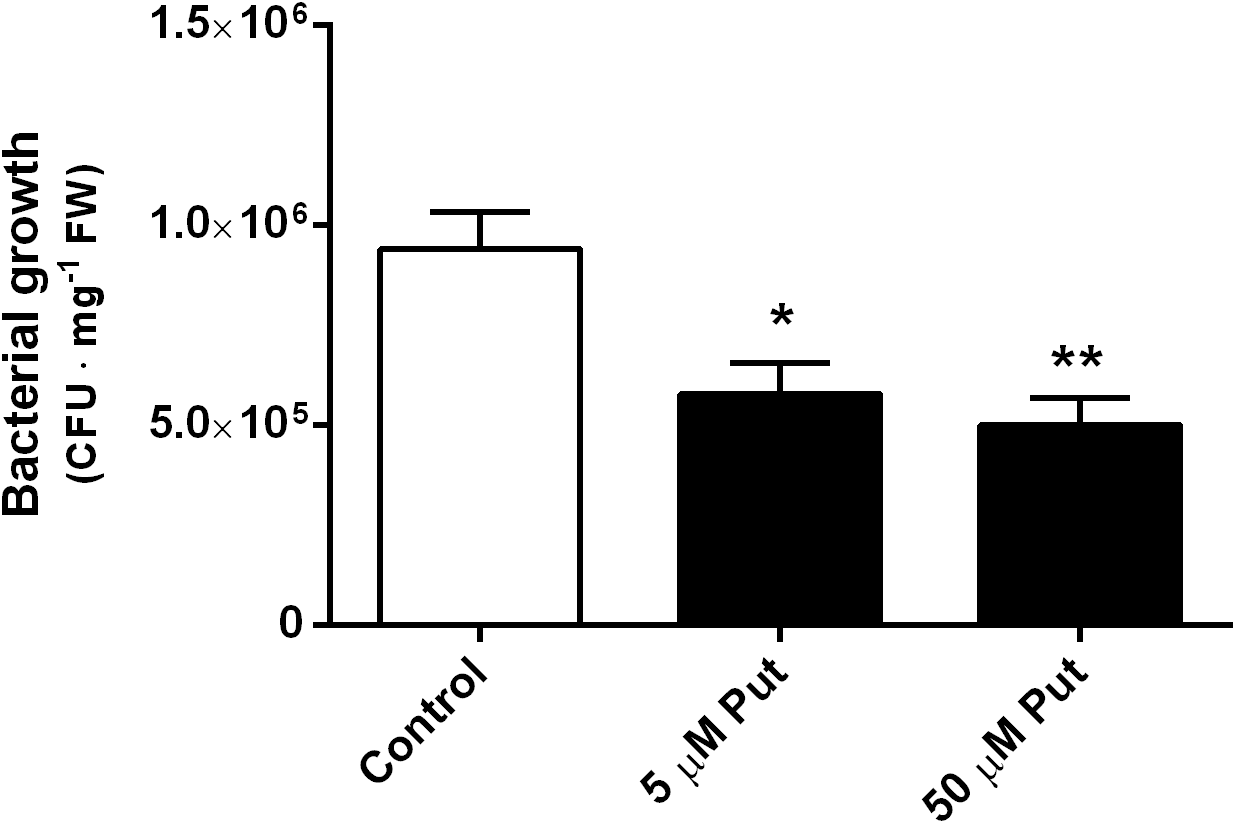
Growth of *P. syringae* pv *tomato* (*Pst* DC3000) in WT Arabidopsis seedlings supplemented with Put. Seedlings were grown in the presence of different Put concentrations and were inoculated by flooding as described in Material and Methods. Seedlings were removed and processed 48 HPI for CFU analysis. Results are means of five replicates ± SEM and statistically significant differences are expressed as *(*P*≤0.05) and **(*P*≤0.01), according to according to one-way ANOVA and Dunnett’s Test

### SA regulates Put biosynthesis and catabolism

In order to evaluate whether the increase in Put mediated by SA depends on *de novo* biosynthesis of this diamine, we assessed the effects of SA on ADC activity, taking into account that this is a key enzyme for Put production in Arabidopsis. As shown in Fig. 4a, treatment of Arabidopsis with 250 μM SA provoked a 40% induction of ADC activity. In addition, we also quantified PA content in two Arabidopsis mutant lines lacking either ADC1 or ADC2 (*adc1-3* and *adc2-3*, respectively), the two ADC isoforms found in this species. In agreement with previous reports (Rossi *et al*., 2015), the concentration of Put in plants growing under control conditions was only compromised in the *adc2-3* line, as Put levels in the *adc1-3* mutant line were similar to those in WT plants (Fig. 4b). These results suggest that ADC2 functions as the main housekeeping enzyme involved in Put homeostasis. In support of this function for ADC2, treatment of *adc1-3* with SA provoked an increase in the concentration of Put, in a similar way to WT plants, while this treatment only caused a very slight increment in Put concentration in *adc2-3* (Fig. 4b). In turn, Spd and Spm levels were less affected than those of Put by SA in WT and *adc* mutants. Thus, whereas SA-treatment provoked a small increment in the concentration of Spd in WT and mutant lines, Spm was only reduced in *adc1-3* and *adc2-3* mutants (Fig. S3).

**Figure 4.**
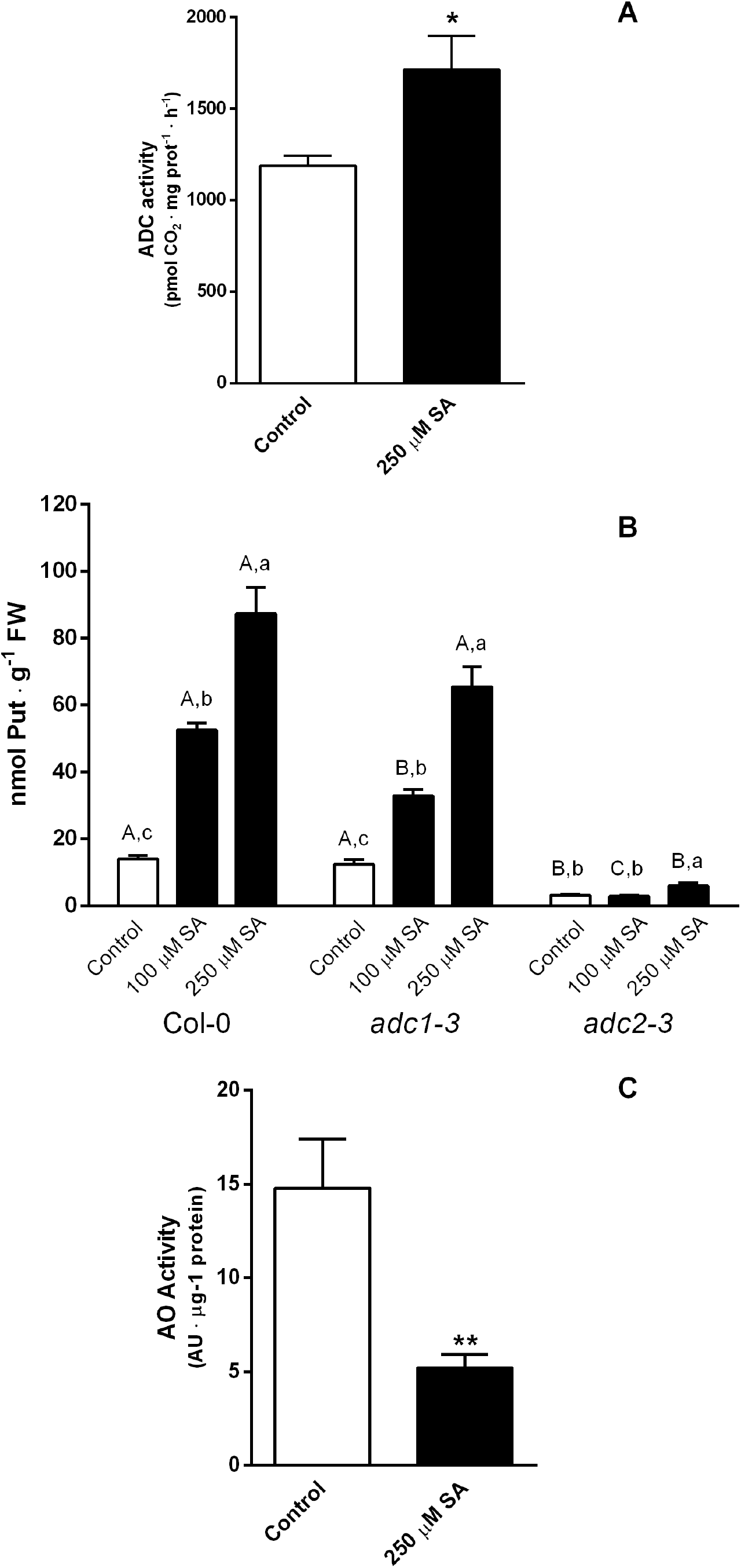
Changes in Put metabolism induced by SA in Arabidopsis seedlings. **A-** Changes in ADC activity of WT *A. thaliana* seedlings in response to SA (250 μM). Seedlings were harvested 48 HPT and immediately used for quantification of ADC activity, which was measured on the basis of the release of ^14^CO_2_ using L-[U-^14^C] arginine as a substrate. Results are means of five replicates ± SEM and statistically significant differences are expressed as *(*P*≤0.05), according to Student’s t-test. **B-** Free Put levels in extracts of WT and *adc*-mutant *A. thaliana* seedlings treated with SA. Polyamines were extracted from control (white bars) and SA-supplemented seedlings 100 μM and 250 μM SA at 48 HPT. Free Put was determined by HPLC after derivatization with dansyl chloride. Results are means of 4-5 replicates ± SEM and different letters indicate statistically significant differences (*P*≤ 0.05) according to twoway ANOVA and Tukey’s multiple comparison test. Capital letters indicate statistically significant differences between plant genotypes within the same treatment. Lowercase letters indicate statistically significant differences between treatments within the same plant genotype. **C-** *In vitro* determination of aminooxidase (AO) activity in AWFs of WT Arabidopsis seedlings. AO activity was determined in AWFs obtained 48 HPT from control and SA-supplemented seedlings (250 μM). For methodological details see materials and methods. Results are means of five replicates ± SEM and statistically significant differences are expressed as **(*P*≤0.01), according to Student’s t-test.

PA levels are not only modulated through *de novo* biosynthesis, but also by means of catabolic enzymes (Moschou *et al*., 2012; Wang *et al*., 2019). Besides its role in the regulation of PA levels, catabolic enzymes are also important because of their contribution to H2O2 production. Put is oxidized by CuAOs, which occur at high levels in dicots. The genome of Arabidopsis contains ten putative CuAO genes, some of which have been well characterized (Tavladoraki *et al*., 2016). Importantly, *AtAO1* and *AtCuAO1* code for apoplastic proteins that utilize Put as the main substrate, even though they are also able to oxidize Spd in some degree (Planas-Portell *et al*., 2013). Therefore, taking into account the increase in apoplastic Put levels induced by SA treatment and the ability of CuAOs to modulate PA levels in the apoplast, we evaluated whether exogenously added SA affects CuAO activity in AWFs of *A. thaliana*. As shown in Fig. 4c, AWFs obtained from SA-treated plants exhibited a 66% reduction in CuAO activity compared to control plants, demonstrating that Put accumulation in the apoplast could also be explained by the inhibition of its catabolism.

### PA metabolism genes are regulated by SA

The results described above demonstrate that SA regulates PA levels through the modulation of enzymatic activities involved in PA biosynthesis and oxidation. We next assessed whether this effect was due to the existence of a mechanism regulating the expression of the cognate genes. For this purpose, we evaluated the expression of the Put biosynthetic genes *ADC1* and *ADC2*, but also *SAMDC1, SAMDC2, SPDS1, SPDS2, SPMS* (all involved in Spd and Spm synthesis) as well as *ACL5* (coding for an enzyme producing the Spm isomer thermo-Spm) (Fig. 5a). In addition, we also included in this analysis the genes encoding for the diamine oxidases *AtAO1* and *AtCuAO1*, and the polyamine oxidases *PAO1, PAO2, PAO3 and PAO4* (Fig. 5b). As a result, exogenous SA induced the expression of most of the PA biosynthetic genes 24 HPT, except for *SPDS2* and *ACL5. ADC2* and *SPMS* up-regulation persisted up to 48 HPT (Fig. 5a). In turn, an induction of the higher PA catabolic genes *PAO1, PAO3* and *PAO4* was observed 24 HPT, the latter still being up-regulated 48 HPT. None of the genes coding for apoplastic Put amine oxidases (*AtAO1* and *AtCuAO1*) showed changes in their transcriptional levels (Fig. 5b).

**Figure 5.**
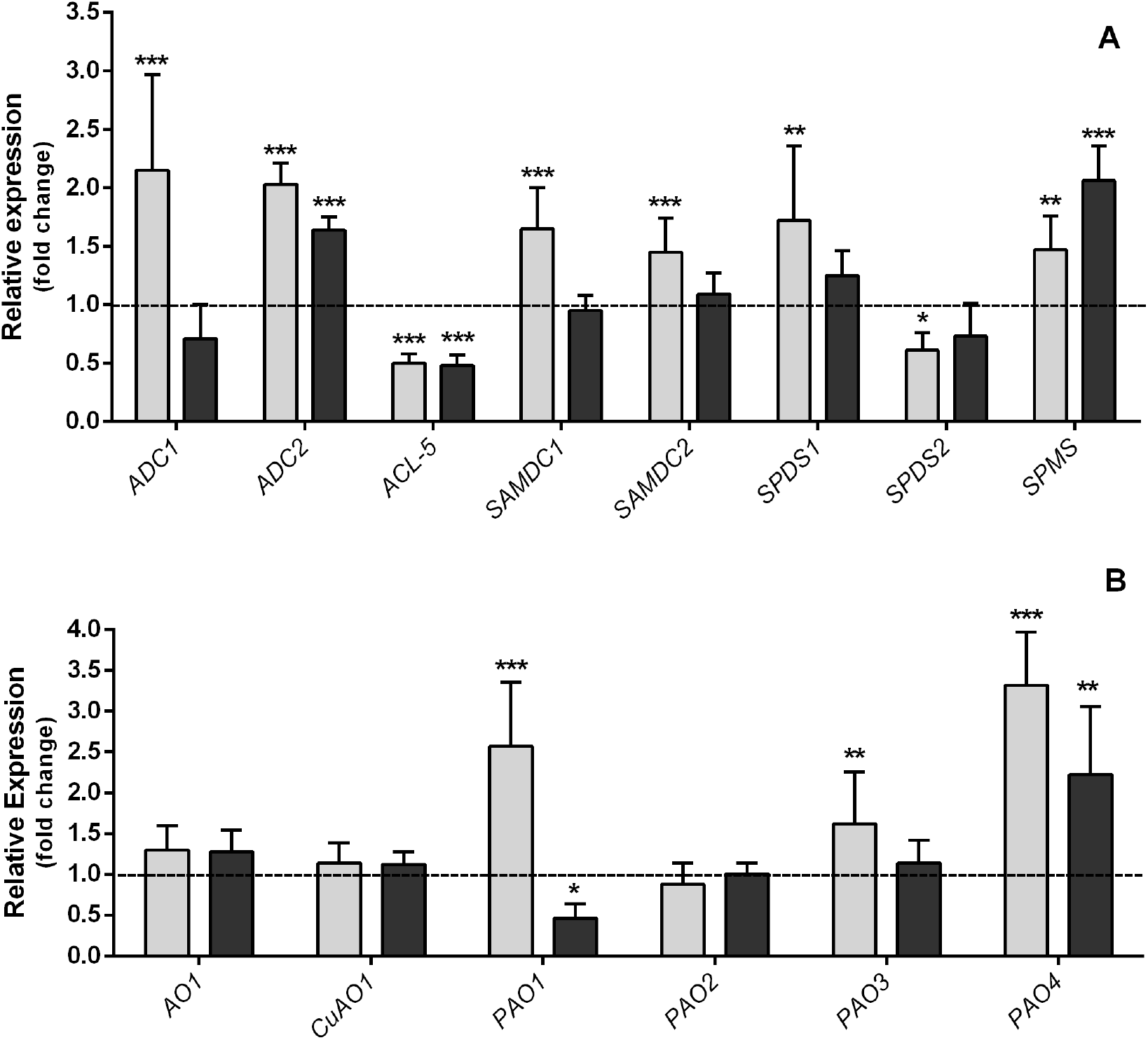
Transcript levels of genes involved in PA synthesis (**A**) and catabolism (**B**) in WT Arabidopsis seedlings supplemented with SA. qRT-PCR was used to analyze the abundance of transcripts in WT seedlings supplemented with 250 μM SA for 24 (gray bars) and 48 (black bars) h. Results are expressed relative to *UBQ10* expression and normalized with respect to control plants, which were assigned to 1 and are represented by a horizontal dotted line. Results presented in both panels are means ± SD of five replicates and statistically significant differences in gene expression, as analyzed with the REST software, are shown as: * (*P*≤0.05), ** (*P*≤0.01) and *** (*P*≤0.001).

### Components of SA signaling involved in the regulation of Put accumulation

Most of the responses associated to SA perception are mediated by the NPR1 protein, which is released from its inactive oligomeric form through cellular redox changes, mainly upon a rise in the cytoplasmic concentration of H2O2 (Fu *et al*., 2012). In addition, other known mediators of the SA response are mitogen-activated protein kinases such as MKK4, MPK6 and MPK3 (Pitzschke *et al*., 2009). Hence, we assessed the contribution of these signaling proteins by evaluating the changes in the concentration of PAs in cognate mutant lines upon SA treatment. Our results show that the accumulation of Put induced by SA is independent of NPR1, as this PA reached similar levels in the *npr1-2* line and the WT after treatment with this hormone (Fig. 6a). SA-induced Put accumulation was not affected in *mkk4* and *mpk3-2* lines, which suggests that MKK4 and MPK3 are not involved in regulating Put accumulation induced by SA. On the contrary, the increase in Put accumulation triggered by SA in the *mpk6-2* mutant was much lower than in WT plants (Fig. 6b), thus evidencing a role for MPK6 in the regulation of PA metabolism by SA-signaling. Anyway, the possibility that MKK4 and MPK3 also participate in the regulation of Put metabolism cannot be completely ruled-out, because of potential functional redundancy with other kinases that could mask the phenotype of *mkk4* and *mpk3-2* mutants.

**Figure 6.**
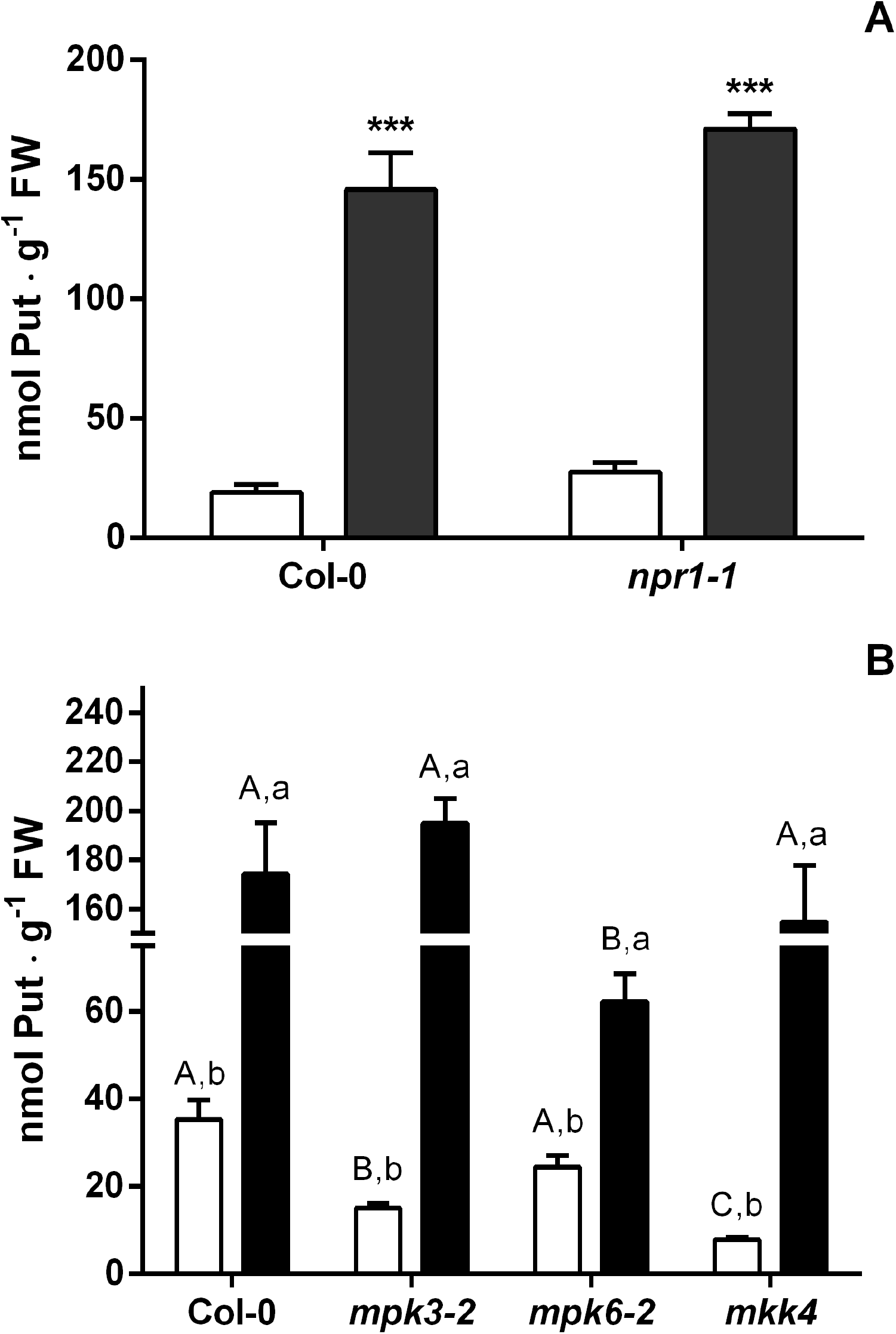
Free Put levels in Arabidopsis mutants of SA signaling proteins. **A-** Changes in Put levels in WT and *npr1-2* mutant Arabidopsis. Fifteen-day-old Arabidopsis seedlings were supplemented with 250 μM SA, 48 HPT control (white bars) and SA-treated samples (black bars) were harvested and processed. Results are means of five replicates ± SEM and statistically significant differences are expressed as *** (*P*≤0.001), according to Student’s t-test. **B-** Changes in Put levels in WT and mitogen-activated protein kinase mutant Arabidopsis. Black bars represent samples obtained from seedlings supplemented with 250 μM SA for 48 HPT, white bars correspond to controls. Free Put was determined by HPLC after derivatization with dansyl chloride. Results are means of five replicates ± SEM and different letters indicate statistically significant differences (*P*≤ 0.05) according to two-way ANOVA and Tukey’s multiple comparison test. Capital letters indicate statistically significant differences between plant genotypes within the same treatment. Lowercase letters indicate statistically significant differences between treatments within the same plant genotype.

### *Pst* infection of Arabidopsis plants triggers Put accumulation in a SA-dependent manner

Results obtained in preceding sections demonstrated that SA induced Put accumulation, and also identified specific components of SA signaling involved in the regulation of Put accumulation. Next, we evaluated the role of SA and SA signaling in regulating Put accumulation during Arabidopsis infection by *Pst*. In this regard, Put levels were assessed in SA-deficient *sid2-2* and *mpk6* plants inoculated with *Pst*. Bacterial infection caused an increase in Put levels in *sid2-2* mutants both 12 and 24 HPI, although in a lesser extent than in WT plants (Fig. 7a-b). At longer post-inoculation times (48h), plants from infected mutant lines showed higher Put levels than WT plants (data not shown). Arabidopsis *mpk6-2* mutants also showed a reduction in Put accumulation 12 HPI with Pst, as compared to WT plants, which provides additional support for the participation of SA signaling in regulating Put accumulation (Fig. 7b). However, Put levels in *Pst*-infected mpk6-2 mutants reached similar values to WT plants 24 HPI (Fig. 7b). As discussed in the previous section, this could be due at least in part to functional redundancy of MPK6 and other kinases involved in SA signaling. Moreover, it is important to highlight that SA-synthesis and signaling mutants used in these experiments, show increased susceptibility to *Pst* infection. In this way, Put derived from the higher number of bacterial cells accumulated in these mutants over time (as compared to WT plants) probably contributed to the increase in Put detected in plant tissues, along with Put derived from plant metabolism. In this regard, it has previously been shown that Put derived from *Pst* is accumulated in infected tomato tissues (Vilas *et al*. 2018). Therefore, Put derived from *Pst* could partly counterbalance the lower Put accumulation exhibited by SA synthesis/signaling mutants, thus explaining the results obtained 24 HPI with *mpk6-2*.

**Figure 7.**
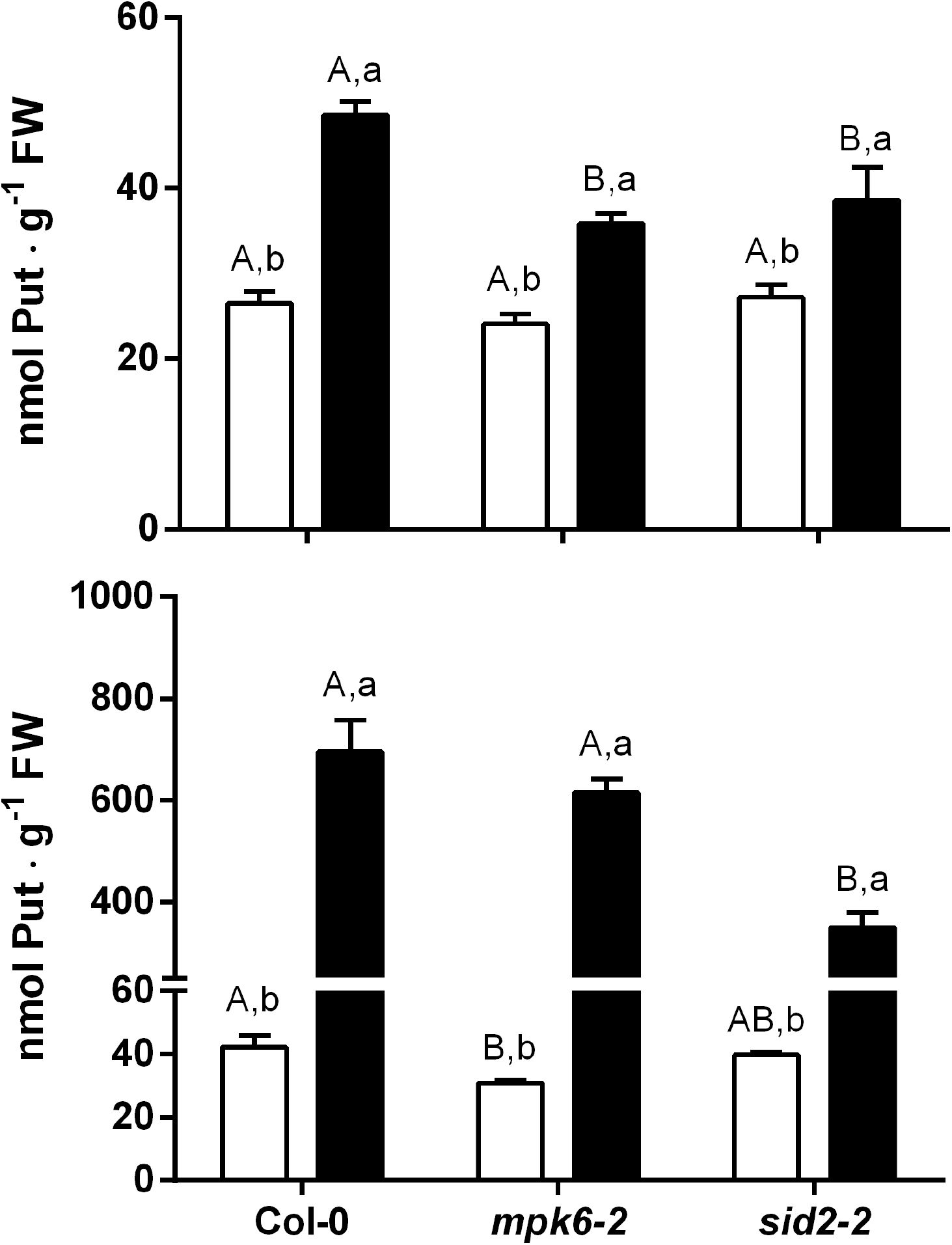
Contribution of the SA pathway to the rise in Put levels after *P. syringae* infection. Put levels in SA synthesis and signaling mutants. Seedlings of WT and *sid2-2* and *mpk6-2* mutant Arabidopsis lines were inoculated by flooding as described in Material and Methods. Seedlings were collected 12 (**A-**) and 24 (**B-**) HPI and processed for PA analysis. Put levels were determined in control (white bars) and inoculated (black bars) seedlings by HPLC after derivatization with dansyl chloride. Results are means of five replicates ± SEM and different letters indicate statistically significant differences (*P*≤ 0.05) according to two-way ANOVA and Tukey’s multiple comparison test. Capital letters indicate statistically significant differences between plant genotypes within the same treatment. Lowercase letters indicate statistically significant differences between treatments within the same plant genotype.

## Discussion

### Modulation of PA metabolism by SA in *Arabidopsis*

SA has been shown to induce PA biosynthesis in several plant species (Németh *et al*., 2002; Jang *et al*., 2009; Zhang *et al*., 2011). Conversely, a recent study by Liu et al (2020) analyzed the role of different plant PAs as inducers of defense signaling pathways mediated by SA in Arabidopsis and demonstrated that Put elicits ROS-dependent activation of SA signaling. Thus, different studies provided evidences about the existence of links between SA signaling and metabolic pathways involved in PA metabolism. However, current knowledge about the mechanisms of SA-mediated regulation of PA metabolism in the context of plant defense is far from being complete. In the present work we used the model plant *A. thaliana* to obtain a detailed picture of the effects of SA on PA metabolism, as well as to analyze the participation of different components of the SA signaling pathway in the regulation of PA metabolism.

In order to explore this issue in the Arabidopsis model, we first determined the SA concentrations able to trigger defense responses in our experimental system without causing detrimental effects on plant growth. Our results demonstrated that 250 μM and even lower SA concentrations increased GUS expression in transgenic *PR1::GUS* plants and also increased plant tolerance to bacterial infection (figure S1 and S2). This is in agreement with previous reports about elevated levels of tolerance to hemibiotrophic bacteria in plants showing higher levels of SA or grown in SA-amended medium (van Wees *et al*., 2000).

PA contents in whole leaf tissues were altered in Arabidopsis plants grow in the presence of SA (figure 1), which depended on the concentration of the hormone and the timing of examination. Particularly, the levels of Put showed the most significant changes, with 6 to 7-fold increments compared to control plants. Accordingly, previous works demonstrated a rise in the concentration of Put in maize and tomato plants in response to SA treatment (Németh *et al*., 2002; Jang *et al*., 2009). The application of the SA-derivative methyl-SA to cherry tomato plants also conduces to the accumulation of Put, Spd and Spm, which was associated to the up-regulation of the biosynthetic genes ADC and ODC (Zhang *et al*., 2011). Similarly, we found that SA-treatment in *Arabidopsis* caused a general induction in the expression of PA biosynthetic genes (figure 5). Importantly, even though the genes encoding for both isoforms of *ADC* (*ADC1* and *ADC2*) were upregulated, the expression of *ADC2* remained induced at the two time-points evaluated. In addition the application of SA to the *adc2-3* null mutant exerted no effect on PA levels, while the *adc1-3* mutant line showed similar levels to WT plants after hormone treatment. Therefore, it seems plausible that ADC2 is the ADC isoform that makes the major contribution to Put accumulation in Arabidopsis during SA signaling. In addition to biosynthesis, Put concentration is also modulated through the regulation of catabolic enzymes. Thus, a rise in the concentration of Put could be explained by a reduction in the rate of its oxidation, or rather stimulation of its production from Spd and Spm by the back-conversion mechanism. Under our experimental conditions, SA up-regulated the expression of three out of four PA oxidase genes assessed (figure 5b), which might explain the reduction in Spm concentration. These results are consistent with the reported stimulatory effects of SA on maize PAO activity, although in this case the effects of this hormone were only observed when applied in association with JA (Angelini *et al*., 2008). Further studies will be necessary to determine if SA stimulates the interconversion of Spm / Spd to Put In Arabidopsis through the action of specific PAO isoforms.

Pathogen recognition leads to considerable changes in metabolic and physiological processes related to cell wall biosynthesis and nutrient transport in the plant apoplast. The concentration of different metabolites like sugars, organic acids, amino acids, secondary metabolites, metals and other cations in this compartment, exert an important impact on the development of plant-microbe interactions (O’Leary *et al*., 2016). For instance, it has been shown that the accumulation and oxidation of PAs in the apoplast is important for defense against a broad range of pathogens in tobacco and Arabidopsis (Marina *et al*., 2008; Romero *et al*., 2018; Liu *et al*., 2019). On this basis, we were prompted to evaluate the effect of SA on PA content in AWFs of Arabidopsis. Our results showed that the profiles of apoplastic PAs are different from those of whole leaf tissues, as Put and Spd were the most abundant PAs. Besides, the addition of SA to the growth medium caused an increment in the concentration of Put, whereas Spd and Spm were slightly diminished (figure 2). Previous works demonstrated the occurrence of a rise in apoplastic Put content during the recognition of virulent and non-virulent microbes (Marina *et al*., 2008; Yoda *et al*., 2009; Vilas *et al*., 2018; Liu *et al*., 2019), suggesting that Put might have an important role in this compartment during the elicitation of plant defense. Even though the rise in Put concentration provoked by SA may be explained by the mechanisms discussed in previous paragraphs, our results also showed that SA treatment leads to a reduction of AO activity (figure 4c). This effect was not associated to a down-regulation in the expression of *AO1* and *CuAO1* (figure 5b), indicating that post-transcriptional mechanisms regulate such activity. Accordingly, it has been shown that SA conduces to a reduction in the activity of CuAO in chickpea plants (Rea *et al*., 2002). On the contrary, Planas-Portel et al. (Planas-Portell *et al*., 2013) demonstrated that SA treatment up-regulates the expression of *AO1* and *CuAO1*. Such discrepancies with the results obtained in the present work could be due to differences in the concentrations and the method used for SA application. Additional work would be necessary to better understand the role of catabolic enzymes in the modulation of apoplastic PA levels by SA.

### MPK6 is involved in the regulation of PA metabolism by SA

SA signaling is mediated by a complex network of regulators, many of which are linked to other hormone signaling systems (Boatwright and Pajerowska-Mukhtar, 2013). As a consequence, SA shows synergistic or antagonistic effects on the responses elicited by other defense hormones. The pathway activated by SA relies mostly on two main components, the regulatory protein NPR1 (Moreau *et al*., 2012) and a set of kinases involving MKK4, MPK3 and MPK6 (Pitzschke *et al*., 2009). In the present work, we used the *npr1-1* line to assess the participation of such protein in the induction of Put accumulation mediated by SA. Our results showed that Put accumulates in the same extension in *npr1-1* and WT plants in response to SA, indicating that the increment in the concentration of this PA is independent of the NPR1 regulator (figure 6a). In this trend, some responses triggered by SA are NPR1-independent (Dong, 2004). In fact, it has been shown that a large part of the genes involved in defense against *Pst* are regulated by NPR1-independent mechanisms (Uquillas *et al*., 2004; Desveaux *et al*., 2004; Blanco *et al*., 2005, 2009). We also assessed the participation of the previously mentioned kinases in the responses associated to SA treatment using mutant lines in the cognate genes. Even though basal Put levels were lower in all three mutant lines included in our experiments (*mpk3-2, mpk6-2* and *mkk4*), only in the *mpk6-2* line Put accumulation was unresponsive to SA-treatment. Therefore, we speculate that MPK6 plays a key role in transducing the signal triggered by SA for modulating Put homeostasis. Arabidopsis MPK6, as well as its tobacco orthologue SIPK, are induced not only by SA but also by other elicitors during pathogen attack. These kinases are crucial for plant tolerance to pathogens, as silencing their cognate genes seriously compromises plant defense responses (Menke *et al*., 2004). The participation of SIPK in the regulation of tobacco PA metabolism has been explored in depth. In this regard, it has been shown that higher activity of NtMEK2 in transgenic tobacco plants, which regulates the activity of SIPK and the alternative kinase WIPK, leads to the upregulation of *ADC*, *ODC* and *PAO* genes, and provokes a dramatic increment of Put concentration (Jang *et al*., 2009). Similarly, induction of MEK2 activity in transgenic *Arabidopsis* lines expressing NtMEK2DD upregulates *ADC1* and *ADC2* and augments Put concentrations, which is suppressed in MPK3 and MPK6 deletion mutants (NtMEK2^DD^/*mpk3* and NtMEK2^DD^/*mpk6*, respectively) (Kim *et al*., 2013).

### SA-mediated Put accumulation during plant responses to *Pst* infection

The present work confirms that inoculation with *Pst* is associated to a rise in Put concentration in *Arabidopsis* (figure 7b), which was previously reported by Kim et al (2013) and Liu et al (2019). This is in agreement with a previous study by our group that demonstrated that *Arabidopsis* infection by a virulent strain of *Pseudomonas viridiflava* causes an increase of Put levels (Rossi *et al*., 2015). Altogether, these results suggest that Put plays an important function in plants during biotic stress. Additional support for a fundamental role of Put in defense responses was provided by a recent study by Liu et al. (Liu *et al*., 2020) In this trend, we show in the present work that Put addition to the culture medium boosts plant immunity (figure 3). On these grounds, we analyzed the participation of the SA pathway in the induction of Put accumulation during *Pst* infection. In this regard, the lower Put accumulation exhibited by the SA-deficient *sid2-2* and the *mpk6-2* mutant shortly after *Pst* inoculation, as compared to WT plants (figure 7), suggests that SA is involved as an early positive modulator of Put accumulation in Arabidopsis plants infected by *Pst*. The higher levels of Put accumulation detected in *sid2-2* and *mpk6-2* mutants at longer times post-inoculation, as compared to WT plants (data not shown) could probably be attributed to the presence of bacterial Put in the PA pool extracted from plant tissues. In this regard, the increased susceptibility of *sid2-2* and *mpk6-2* mutants to bacterial infection is known to lead to higher bacterial titers than in WT plants, which could counterbalance the lower accumulation of plant-derived Put in the mutant lines. Further work would be necessary to test this hypothesis, but a straightforward experimental approach cannot be easily envisaged, because analytical techniques of PA analysis do not allow distinguishing between plant and bacterial Put.

### Conclusions

SA causes a notable increment in Put levels in whole plant extracts of Arabidopsis, but also specifically in apoplastic fluids, mainly due to the induction of ADC2. The rise in the concentration of Put elicited by SA does not depend on the NPR1 regulator, but is positively regulated by MPK6. In this way, SA acts as a regulator of PA metabolism, which could explain the increment in Put levels reported for several plant-pathogen interactions in which plant defenses are mediated by this hormone.

## Abbreviations

ADC: arginine decarboxylase;
AOs: amine oxidases;
AWFs: apoplastic washing fluids;
CFUs: colony forming units;
Col-0: Columbia-0;
CuAO: copper-containing amine oxidase;
HPT: hours post-treatment;
NPR1: Non-Expressor of Pathogenesis Related 1;
ODC: ornithine decarboxylase;
PAO: polyamine oxidase;
PAs: polyamines;
*Pst*: *Pseudomonas syringae* pv. *tomato* strain DC3000;
Put: putrescine;
SA: salicylic acid;
SAMDC: S-adenosylmethionine decarboxylase;
Spd: spermidine;
SPDS: spermidine synthase;
Spm: spermine;
SPMS: spermine synthase.

## Supplementary data

**Figure S1.** *In vitro* GUS activity of *PR1p*::GUS upon SA treatment.

**Figure S2.** Growth of *P. syringae* pv *tomato* (*Pst*) DC3000 in WT Arabidopsis seedlings supplemented with SA.

**Figure S3.** Free PA levels in extracts of WT and *adc* mutant *A. thaliana* seedlings treated with SA.

**Table S1.** List of primers used for quantitative real-time PCR.

## Acknowledgements

FRR, AG, MM and FLP are members of the Research Career of CONICET. The authors are grateful to Patricia Alejandra Uchiya (CIC) for valuable technical assistance. This work was financially supported by grants of Agencia Nacional de Promoción Científica y Tecnológica (PICT 2012-1716 and PICT 2014-3286) and CONICET (PIP 2010-0395 and PIP 2014-0903) to FLP.

## Author contributions

FLP, FRR and MM formulated the research project. FLP obtained funding for supporting the research. FRR performed the experimental work. FRR, AG and FLP analyzed and discussed the results. FRR and FLP wrote the manuscript with the collaboration of AG. All the authors approved the manuscript.

## Data availability statement

The biological materials used in this study area available upon request.

